# Aging and intraocular pressure homeostasis in mice

**DOI:** 10.1101/2023.10.17.562768

**Authors:** Guorong Li, Joseph van Batenburg-Sherwood, Babak N. Safa, Nina Sara Fraticelli Guzmán, Andrea Wilson, Mohammad Reza Bahrani Fard, Kevin Choy, Michael L. De Ieso, J. Serena Cui, Andrew J Feola, Tara Weisz, Megan Kuhn, Cathy Bowes Rickman, Sina Farsiu, C. Ross Ethier, W. Daniel Stamer

## Abstract

Age and elevated intraocular pressure (IOP) are the two primary risk factors for glaucoma, an optic neuropathy that is the leading cause of irreversible blindness. In most people, IOP is tightly regulated over a lifetime by the conventional outflow tissues. However, the mechanistic contributions of age to conventional outflow dysregulation, elevated IOP and glaucoma are unknown. To address this gap in knowledge, we studied how age affects the morphology, biomechanical properties and function of conventional outflow tissues in C57BL/6 mice, which have an outflow system similar to humans. As reported in humans, we observed that IOP in mice was maintained within a tight range over their lifespan. Remarkably, despite a constellation of age-related changes to the conventional outflow tissues that would be expected to hinder aqueous drainage and impair homeostatic function (decreased cellularity, increased pigment accumulation, increased cellular senescence and increased stiffness), outflow facility, a measure of conventional outflow tissue fluid conductivity, was stable with age. We conclude that the murine conventional outflow system has significant functional reserve in healthy eyes. However, these age-related changes, when combined with other underlying factors, such as genetic susceptibility, are expected to increase risk for ocular hypertension and glaucoma.

## 1. Introduction

Age and elevated intraocular pressure (IOP) are the two primary risk factors for glaucoma [1-6], the leading cause of irreversible blindness [7]. The prevalence of glaucoma prevalence increases from 0.2% at age 50 to 2.7% at age 59, and can reach 12.8% in those over 80 [6, 8-10]. With life expectancies increasing, it is predicted that there will be 1.5 billion people worldwide over the age of 65 by 2050 [11]. Hence, a dramatic increase in the incidence of glaucoma is expected in the years to come.

Both the incidence of glaucoma and its progression are associated with ocular hypertension (elevated IOP) [12-20]. While IOP depends on several factors, it is the resistance to aqueous humor outflow that largely sets IOP, and in a healthy human eye, maintains it in the normal range over a lifetime [21-30]. In other words, elevated aqueous outflow resistance causes ocular hypertension [31]. In both normotensive and hypertensive eyes, outflow resistance is (dys)regulated by the coordinated function of the trabecular meshwork (TM), a porous connective tissue, and Schlemm’s canal (SC), an endothelial-lined circular fluid collecting duct in the anterior eye. These two structures (i.e., TM and SC) are the key elements of the so-called conventional outflow pathway, through which the majority of the aqueous humor flows out of the eye (Figure 1).

**Figure 1:**
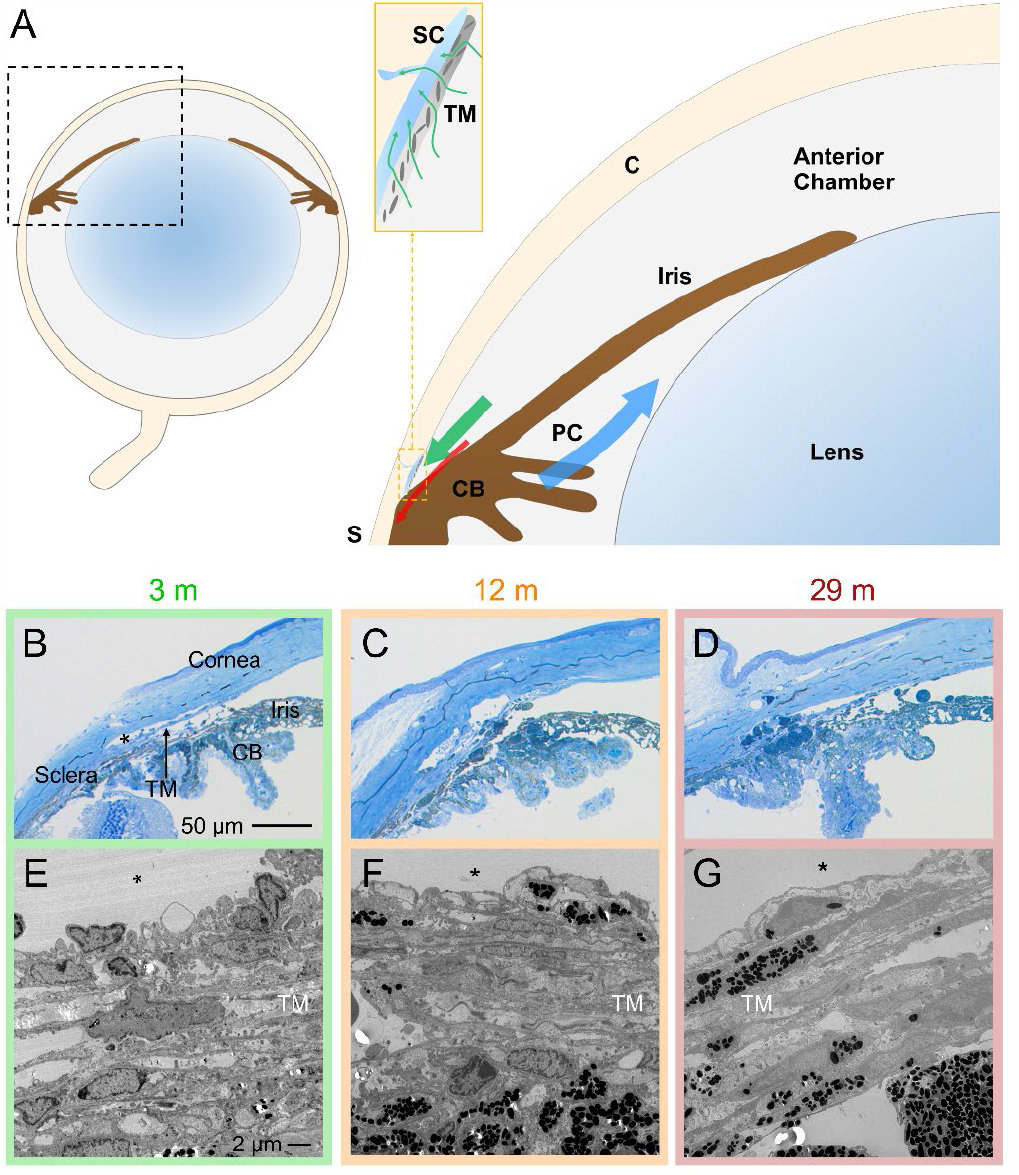
Schematic and morphology of the irideo-corneal angle tissues of the mouse eye. **A)** The blue arrow, located in the posterior chamber (PC, behind the iris), shows the direction of aqueous humor flow after production by the ciliary body (CB). Aqueous humor enters the anterior chamber of the eye through the pupil, and then circulates in the anterior chamber before draining from the eye. The large green arrow shows the direction of aqueous humor flow out of the eye, primarily through the conventional outflow pathway (inset), and secondarily through the unconventional pathway (red arrow). The two key tissues of the conventional outflow pathway are the trabecular meshwork (TM), a porous connective tissue, and Schlemm’s canal (SC), a specialized collecting vessel with an endothelial lining that has characteristics of both lymphatic and vascular endothelia. Additional abbreviations: Cornea (C) and Sclera (S). Below the schematic are representative histologic images (sagittal sections) of anterior segments from three different age groups of mice imaged by light microscopy **(B-D)** or transmission electron microscopy **(E-G)**. While the gross structure of outflow tissues did not change with age, older eyes contained more pigment and pigment-laden cells in the TM. * Denotes Schlemm’s Canal lumen; images shown are representative, and are from mice of 3, 12 and 29 months of age.

Despite most individuals having IOP in the normal range, age-related changes occur in the human conventional outflow pathway in both normal and glaucomatous eyes. For example, there is a steady loss of cells in the conventional outflow pathway with age that is accelerated in glaucoma [32, 33]. Matching a decrease in aqueous inflow (production) rate [20], both unconventional and conventional outflow drainage rates decrease with age to normalize IOP [34, 35]. Aging is an inevitable and complex multifactorial biological process that predisposes tissues to disease [36, 37]. To better understand the complex processes that contribute to ocular hypertension and glaucoma, it is essential to know more about age-related changes in the conventional outflow tissues.

To study these complex issues, we used mice, for several reasons. Mice are widely used to model human disease, including glaucoma. Importantly, conventional outflow tissues in mice are anatomically, physiologically and pharmacologically remarkably similar to humans [38-42]. Further, mice have a much shorter lifespan than humans, facilitating aging studies. Finally, the thin murine sclera allows *in vivo* imaging of outflow pathway tissues [41, 43]. Therefore, we designed a study to monitor age-related changes in the conventional outflow pathway in mice. Using mice that ranged in ages from 2 to 32 months (equivalent to 16-to 80-year-old humans), we examined conventional outflow tissue morphology, biomechanical properties and function.

## 2. Materials and Methods

### Animals

A total of 213 C57BL/6J (C57) mice (both males and females, ages 2-32 months) were used in this study for the various measurements performed. It was not possible to conduct all reported measurements on each mouse (histology, OCT imaging, perfusion), and thus each type of experiment was conducted on a subset of the 213 mice, so that age ranges reported in individual graphs and tables may not span the entire range from 2-32 months. All mice were handled in accordance with approved protocols (A001-19-01 and A226-21-11; Duke University Institutional Animal Care and Use Committee) and in compliance with the Association for Research in Vision and Ophthalmology (ARVO) Statement for the Use of Animals in Ophthalmic and Vision Research. The mice were purchased from the Jackson Laboratory (Bar Harbor, Maine, USA), bred/housed in clear cages and kept in housing rooms at 21°C with a 12h:12h light:dark cycle.

### Body mass measurements

An empty cage was tared on a balance (Model 1015SSTP, Taylor) after which a live mouse was put into the cage and the body mass was recorded.

### Intraocular pressure measurements

The mice were anesthetized with intraperitoneal injected ketamine (60 mg/kg) and xylazine (6 mg/kg). IOP was measured immediately upon cessation of movement (i.e., under light anesthesia) using rebound tonometry (TonoLab, Icare, Raleigh, NC, USA) between 10am and 1pm [38, 40, 41, 68, 69]. Each recorded IOP was an average of six measurements, giving a total of 36 rebounds from the same eye per recorded IOP value.

### Perfusion measurement of aqueous humor dynamics parameters

We measured key parameters of aqueous humor dynamics (AHD), namely outflow facility, β (nonlinearity factor) and ocular compliance in freshly enucleated mouse eyes using the iPerfusion system as described in detail previously [38, 39, 48, 70, 71]. Briefly, mice were euthanized using isoflurane gas, and eyes were carefully enucleated. Each eye was mounted on a stabilization platform in the center of a temperature-regulated perfusion chamber using a small amount of cyanoacrylate glue (Loctite, Westlake, OH, USA). The perfusion chambers were filled with pre-warmed Dulbecco’s phosphate-buffered saline with added D-glucose (DBG 5.5 mM), submerging the eyes to maintain their hydration, and temperature at 35°C. The eyes were then cannulated using glass microneedles connected to each channel of the iPerfusion system. Using micromanipulators under a stereomicroscope, a microneedle was inserted into the anterior chamber of each eye. The eyes were perfused at 9 mmHg for 30 min to allow acclimatization and stabilization, followed by perfusion at 9 sequential pressure steps of 4.5, 6, 7.5, 9, 10.5, 12, 15, 18 and 21 mmHg. To define stability at each pressure step, the flow rate (*Q*) was filtered with a 60-second Savitsky-Golay filter [72], and its time derivative (the slope) was determined via linear regression analysis. Stability was defined as a minute-long maintenance of the slope under 3 nl/min/min. Stable *Q* and pressure (*P*) data were filtered with a 60-second Savitsky-Golay filter and averaged over 4 minutes at each pressure step for further analysis [40, 48, 69].

Outflow facility and *β* and their 95% confidence intervals were calculated using non-linear regression to fit the established power law model to the experimental data

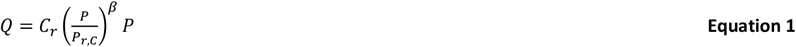

where *C*_*r*_ is the ‘reference facility’ measured at the reference pressure, *P*_*r*_,*C* = 8 mmHg, chosen to represent a physiological pressure drop across the conventional outflow pathway [48].

To calculate the ocular compliance, defined as the instantaneous change in the eye volume in response to a unit change in IOP, we employed the step response method, described in detail previously [71].

Briefly, a lumped parameter model was generated of the components of the perfusion system and eye, with facility as in Equation 1 and ocular compliance given by

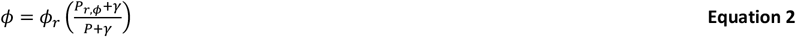

where *ϕ*_*r*_ is the ‘reference compliance’ measured at a reference pressure, *P*_*r*_,*ϕ* = 13 mmHg (chosen for consistency with previous studies [71]. The term *γ* is an arbitrary fitting parameter. The model yields an equation for the change in intraocular pressure in response to a step increase in perfusion system pressure, which can be fit to the measured pressure data to calculate *ϕ*(*P*). Finally Equation 2 can be fit to the measured *ϕ*(*P*) values using non-linear regression to extract a value for *ϕ*_*r*_ [71].

### Ocular diameter measurements

Since our measured AHD parameters can vary with eye size, we measured the diameter of a subset of eyes to adjust for that variability (Supplementary Figure 1). In brief, enucleated eyes were cleaned by carefully dissecting away conjunctiva and extraocular muscles. Eyes were then placed on a ruler (as a reference scale) and imaged under a dissecting microscope. The images were analyzed using ImageJ [73] software in a masked fashion, with the diameter at the ocular equator measured in millimeters.

Ocular diameter was only measured for a subset of the eyes, leading to three groups:

- Group I: Eyes in which diameter was measured but AHD parameters were not,
- Group II: Eyes in which AHD parameters were measured, but ocular diameter was not, and
- Group III: Eyes in which both diameter and AHD parameters were measured.

To enable the adjustment of measured AHD parameters for ocular diameter in Group II eyes, we estimated the diameter of eyes for which the data was missing using non-linear regression. We found empirically (Figure S1B) that the plot of diameter vs. age (for Groups I and III) was reasonably fit by

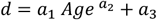

Where *a*_1_ [mm/month], *a*_2_ [-] and *a*_3_[mm] were the fitting parameters. Diameters for eyes for which *d* was not available were then estimated using the best-fit values (Figure S1C).

### Outliers and sex differences

We first filtered AHD data for outliers, performed on a per-age-group basis for the logarithms of both facility and compliance, since both quantities are known to be log-normally distributed [48, 71]. We eliminated any data points that were more than 2.5 median absolute deviations away from the median, accounting for the 95% confidence interval on the parameters [48]. This led to a single eye in the oldest group being rejected based on high compliance. Data were not adjusted or stratified for sex because we did not observe a statistical difference in eye size between males and females of the same age.

### Adjustment of ocular compliance and outflow facility to account for eye size

Facility is expected to be dependent on the cross-sectional area of the inner wall of SC through which the fluid filters. This area is equal to the circumference of the toroidal SC (approximately equal to limbal circumference) multiplied by SC anterior-posterior length, both of which have been observed to increase with age, but only modestly [64]. Normalization by area would give a conductivity (facility/area) with hard-to-interpret units [nl/min/mmHg/mm^2^] and involve an estimate of SC anterior-posterior length, which showed significant inter-animal variability [64]. Instead, we used our correlation between age and ocular diameter to adjust the measured reference facility to the value that would be expected for an eye of the average diameter of the eyes used in this study (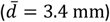). This “adjusted reference outflow facility”,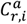, for a given eye, *i*, is given by

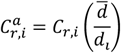

The value 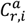 is the adjusted facility of the eye if it were to have diameter 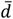 rather than its actual diameter *d*_*i*_. We plot the unadjusted facility, *C*_*r,i*_, in Figure 5B, which is the physiologically relevant quantity strongly influencing IOP, while the adjusted value, 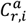, is plotted in Supplemental Figure 3A and as black dots in Figure 5B.

Ocular compliance is, by definition, dependent on eye volume (*πd*^3^/6). Similar to facility, we used ocular diameter to adjust the measured compliance to the compliance that the eye would have had if it had diameter 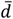

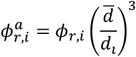

where 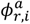 is the adjusted compliance of eye *i*, and *ϕ*_*r,i*_ is the compliance of eye *i* before adjustment. Figure 6B plots 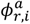, which is a relevant measure of ocular coat (corneo-scleral shell) material properties, while Supplemental Figure 3B plots the unadjusted compliance values *ϕ*_*r,i*_.

Note that since compliance and facility are both log-normally distributed, we carried out all analyses of these quantities in the logarithmic domain [48, 71].

### Optical coherence tomographic imaging

*In vivo* imaging was carried out using an Envisu R2200 high-resolution spectral domain (SD)-OCT system (Bioptigen Inc., Research Triangle Park, NC, USA). We followed our previously established techniques to image iridocorneal angle structures in mice [38-41, 68, 70, 74]. Briefly, mice were anesthetized with ketamine (100 mg/kg)/xylazine (10 mg/kg) by IP administration. While mice were secured on a custom-made platform, a single pulled glass microneedle filled with phosphate buffered saline (PBS) was inserted into the anterior chamber of one eye. The microneedle was connected to both a manometric column to adjust IOP and a pressure transducer to continuously monitor IOP levels using PowerLab software. The OCT imaging probe was aimed at either the nasal or temporal limbus, and the image was centered and focused on the SC lumen. While collecting images, mouse eyes were subjected to a series of IOP steps (10, 12, 15, 17 and 20 mmHg) by adjusting the height of the fluid reservoir. At each IOP step, a sequence of repeated OCT B-scans (each with 1000 A-scans spanning 0.5 mm in lateral length) from spatially close positions were captured, registered and averaged to create a high signal-to-noise-ratio image of the iridocorneal angle region in each animal. The duration of each pressure step was ∼1-2 minutes.

### Segmentation of OCT images

OCT B-scans of iridocorneal angle tissues were registered and segmented following established methods [38-40] using SchlemmSeg software, which includes two modules: Schlemm I and Schlemm II. Briefly, OCT B-scans were automatically registered using our custom Schlemm I software for SC segmentation. The Schlemm II software package was then used to differentiate SC from scleral vessels, which were automatically marked. If SC was seen to be connected to collector channels (CC), manual separation of SC from CC was required, and was based on the shape of SC and speckling in the images generated by blood cells or other reflectors contained in blood vessels [39-41, 70, 75-78]. The speckle variance OCT-angiography images were generated based on the speckling in SC and vessels as described in detail previously [40]. SC was easily differentiated from other vessels due to its large lumen size and location.

### Measurement of iris-cornea subtended angle

It was evident from OCT scans that the iris bowed and displaced posteriorly as IOP increased. To quantify this effect, we measured the angle subtended by the iris and cornea. In anesthetized mice (see above), OCT images of the anterior segment (400 × 300 pixels, corresponding to a 2 × 1.26 mm field of view [horizontal × vertical]) were imported into ImageJ, and 2-line segments, each 19 pixels long, were drawn from the apex of the irideo-corneal angle to the anterior surface of the iris and to the posterior surface of the cornea. The angle subtended between these segments was measured in ImageJ, accounting for the horizontal-to-vertical aspect ratio of the image.

### Quantitative estimation of TM biomechanical properties (tissue stiffness)

The stiffness of the TM increases in glaucomatous eyes and is known to be associated with the function of the conventional outflow pathway tissues, i.e., stiffer tissues have higher flow resistance [45]. It was therefore of great interest to assess how TM stiffness changes with age. For this purpose, we used finite element modeling driven by OCT images acquired as described above. In brief, increasing IOP deforms the TM and partially collapses SC lumen; we tracked luminal collapse as the main OCT imaging outcome measure, which we then fit with our finite element model. The full modeling methodology is beyond the scope of this paper and is described elsewhere [64]. In brief, we used the FEBio finite element software [80] to simulate age-specific models, evaluating TM/SC deformations vs. IOP and adjusted TM stiffness so that finite element predictions of SC lumen partial collapse matched those observed experimentally by OCT. This approach included the effects of posterior deformations of the iris, pectinate ligaments connecting the iris and TM, and changes in posterior chamber pressure and SC luminal pressure with IOP. At IOPs of 15 and 20 mmHg, we computed a difference in normalized SC area, Δ*nSCLA*, as follows.

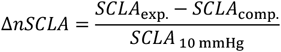

where *SCLA* is the cross-sectional area of Schlemm’s canal lumen, the subscript “comp.” refers to a computed result (finite element simulation), the subscript “exp.” Refers to an experimental result (OCT imaging), and the subscript “10 mmHg” refers to the reference value of *SCLA* at an IOP of 10 mmHg. Δ*nSCLA* is essentially a measure of how closely the computed collapse of SC lumen agrees with the experimentally observed collapse as IOP is increased. Perfect agreement corresponds to Δ*nSCLA* = 0.

### Histology and transmission electron microscopy

Mouse eyes were carefully enucleated immediately after sacrifice as described above, and immersion-fixed in 4% paraformaldehyde at 4 °C overnight. The eyes were then bisected, and the posterior segments and lenses were removed. The anterior segments were cut into four quadrants.

For immunohistochemistry studies, one quadrant was embedded into OCT and 10 µm sections were cryosectioned and immunostained with antibody against p21 (1:500 dilution, rabbit polyclonal, ab188224; Abcam). The secondary antibody was Alexa Fluor® 594 AffiniPure Goat Anti-Rabbit IgG (H+L) (Jackson ImmunoResearch Laboratories INC) at 1:500 dilution. Nuclei were stained with DAPI (1:1000), and images were captured using a Nikon Eclipse 90i confocal laser-scanning microscope (Melville). Images were collected at identical intensity and gain settings for all sections. Using ImageJ software, the TM from each image was manually segmented (region of interest, ROI) in a masked fashion (by MK), and red fluorescence intensity (mean grayscale value) within each ROI was determined. The inner wall of SC provided the outer boundary and the iris/TM interface the inner boundary of the ROI. Only DAPI-labeled areas in TM regions were selected for analysis.

For histology studies, two opposite quadrants per eye were embedded in Epon, and 0.5 µm semithin sections were cut, stained with 1% methylene blue, and examined by light microscopy (Axioplan2, Carl Zeiss MicroImaging, Thornwood, NY).

For electron microscopy studies, mouse anterior quadrants were embedded in Epon resin and 65 nm sagittal ultrathin sections were cut through iridocorneal angle tissues using an ultramicrotome (LEICA EM UC6, Leica Mikrosysteme GmbH, A-1170, Wien, Austria). Sections were stained with uranyl acetate/lead citrate and examined with a JEM-1400 electron microscope (JEOL USA, Peabody, MA). In each section, cell numbers (nuclei) and cells containing engulfed pigment were counted in the entire visible TM tissue area (by JSC or GL) and pigment granules were counted within 7 µm of the inner wall of SC (by AW) in a masked fashion. The counts were normalized by the visible TM area and are presented as number of cells per mm^2^ of entire TM or number of pigment particles per mm^2^ of TM within 7 μm of the inner wall of SC.

### Statistical analysis

We evaluated whether the AHD parameters (IOP, facility, beta and compliance) measured in paired eyes were correlated (Supplemental Figure 4). Although facility was not correlated (p > 0.05), pairwise correlation coefficients for IOP, beta and compliance were greater than 0.5 with p < 0.001; thus, measurements from individual eyes of a pair could not be treated as independent. Therefore, the bilateral averages from each pair of eyes were used to represent each animal and the confidence intervals for facility, beta and compliance were added in quadrature. There were some mice in which only one eye perfusion was successful; values measured in these eyes were included without modification (open circles in Figures 3B, 7A and 7B).

To investigate the relative change in SC area with pressure (Figure 4 A-C), linear regression with a fixed intercept at 100% was used to calculate the change in % SC area/mmHg and the corresponding 95% confidence interval for each eye (Figure 4D).

We also observed that the iris was displaced posteriorly as IOP increased, and the extent of this displacement was age-dependent. To characterize this phenomenon, we used OCT images to measure the angle subtended by the cornea and the root of the iris as a function of intraocular pressure. The increase in angle from baseline (*θ* − *θ*_0_) was compared to the logarithm of the increase in pressure from baseline (*P* − *p*_0_) using an empirical fit of the form

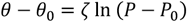

where *ζ* quantifies the sensitivity of irideo-corneal angle changes to pressure and the subscript “0” refers to the baseline condition of IOP = 10 mmHg.

Although three broad age groups were considered, age was treated as continuous variable. Regression on age was used to calculate coefficients of determination (R^2^) for cellularity, pigment granule density, IOP, log-transformed adjusted reference outflow facility, change in SC area per mmHg, baseline irideocorneal angle, sensitivity of irideo-corneal angle to pressure (ζ), *β* and log-transformed adjusted reference ocular compliance. Regression parameters are reported as mean [95% confidence interval].

When testing compliance, outflow facility, and *β*, we used Bonferroni correction to adjust p-values in order to avoid increasing the risk of a type I error since three comparisons were made on the same data. Thus, for these parameters, we report *p*_*B*_ = 3*p*. All p-values were considered relative to α = 0.05, 0.01 or 0.001.

## 3. Results

### Eye size increased with age

We measured globe size and body mass in mice of different ages. As expected, both increased with age (Supplementary Figure 1). Interestingly, globe size increased more dramatically at early stages of life than later for both sexes. Body mass of both males and females steadily increased throughout their lifetime (Supplementary Figure 1).

### Aging reduces TM cellularity, but increases pigment and levels of the senescence marker CDKN1A (p21) in the TM

To determine whether outflow tissue structure changes over time, anterior segments of C57 mice at different ages (from 2 to 32 months old) were processed for morphological evaluation by both light and electron microscopy (Figure 1). Mice were grouped into 3 age cohorts: young (<8 months old), middleaged (8-16 months old) and elderly (>16 months old). At the light microscopic level (Figure 1 B-D), the gross appearance of the conventional outflow tissues (i.e., TM beam architecture and shape of SC lumen) did not differ appreciably between the three age groups. However, we did observe greater amounts of pigment and greater numbers of pigment-laden cells in the TM of middle-aged and elderly mice vs. young mice.

To quantify these morphological changes, we examined transmission electron micrographs (Figure 1 E-G). Treating age as a continuous variable, we quantified the number of cells in the TM and observed a steady and significant decline in cell density of -2.0 [-2.8,-1.3] cells/mm^2^/month (mean [95% confidence interval (CI)]; *R*^2^ = 0.66, *p* < 10^−3^; Figure 2A). In the same samples, we also counted the number of pigment granules in the juxtacanalicular tissue (JCT; here defined as being within 7 µm of the inner wall of SC), observing an age-dependent increase in pigment granule density of 6.8 [1.7, 11.9] pigment granules/mm^2^/month (*R*^2^ = 0.32, *p* < 0.05; Figure 2B). Further, the cells remaining within the TM displayed an age-related increase in the senescence marker CDKN1A (p21, Figure 2 D-I). Evaluation of the intensity of the fluorescent signal indicated a clear increase in expression of p21 with age (R^2^=0.78, p<0.001, Figure 2C).

**Figure 2.**
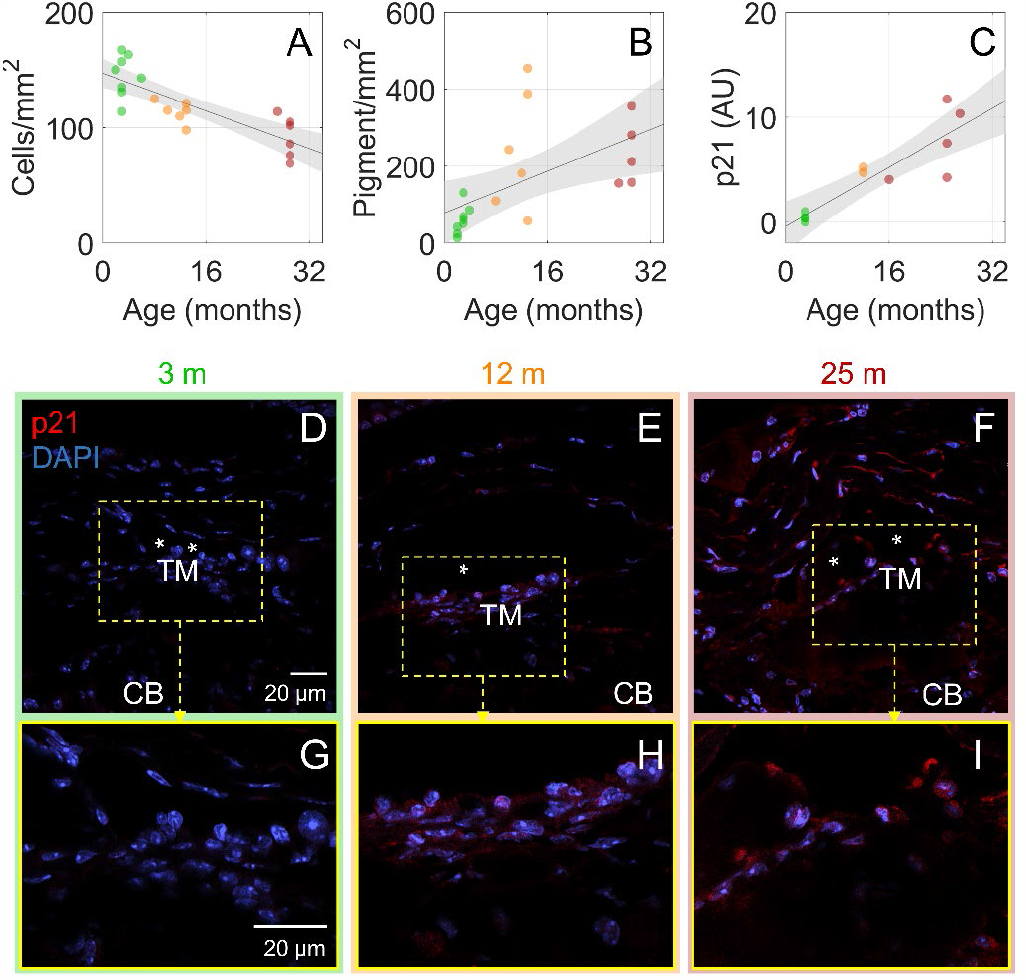
Decreased cellularity, increased pigment, and increased p21 expression were observed in the trabecular meshwork of older mouse eyes. **A)** The number of cell nuclei in the TM were counted in transmission electron micrographs (600×) and normalized by the visible TM area. An age-associated reduction in cellular density was evident (R^2^ = 0.66, p < 0.001). **B)** The number of pigment granules within 7 µm of the inner wall of SC was counted and normalized to TM area. The density of pigment granules was weakly correlated with age (R^2^ =0.32, p < 0.05). **C)** The intensity of the fluorescent signal from p21 labeling of cells in the conventional outflow pathway from three age groups of mice (young, middle-aged and elderly) was found to significantly increase with age (R^2^=0.78, p<0.001). Lines show best linear fits with grey shaded regions indicating the 95% confidence bands. Colors: green – young, orange – middle-aged, and red – elderly. **D-F)** p21 (red) and nuclei (DAPI, blue) labeling from sagittal sections of outflow tissues taken at the same confocal microscopic settings. The bottom row shows an enlargement of the region in the dashed box in the top row. * Denotes the lumen of SC; CB, ciliary body. Images are representative of N = 3-4 mice/age group.

### TM tissue stiffens with age

We next investigated whether these age-associated changes in the TM were accompanied by changes in TM biomechanical properties, since changes in such properties are known to occur in ocular hypertension and are associated with TM dysfunction [38, 39, 44-47]. Using high-resolution spectral-domain optical coherence tomography (OCT), we monitored deformations in conventional outflow tissues in response to increasing IOP. Since IOP-mediated changes in SC lumen dimensions depend upon TM stiffness [38, 39], we specifically monitored SC lumen dimensions over five sequential IOP steps (see sample images at 3 of these steps in Figure 3), using our custom semi-automatic segmentation algorithm to identify SC lumen from OCT images.

**Figure 3.**
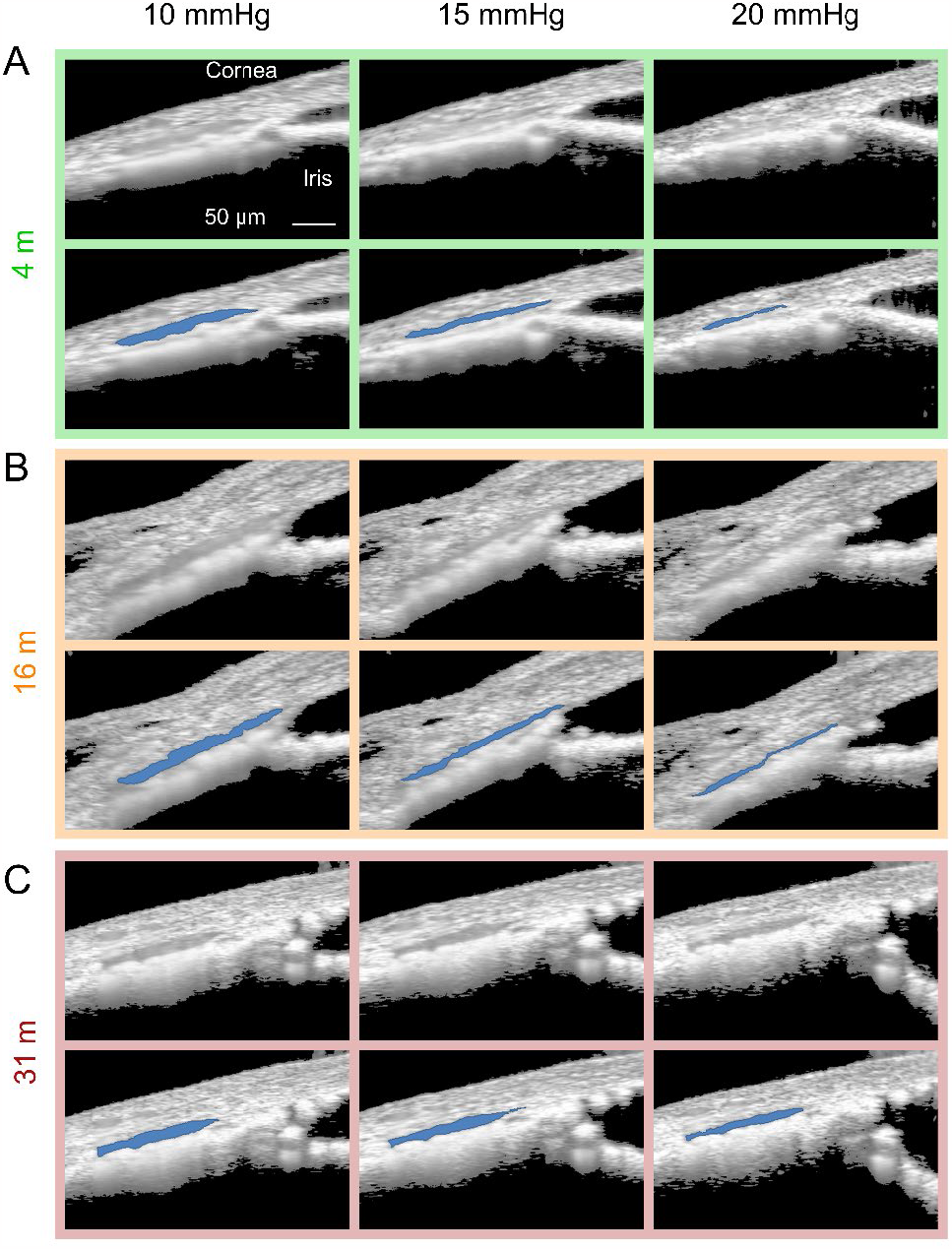
Schlemm’s canal is less susceptible to IOP-induced collapse in older mice. The response of conventional outflow tissues to sequentially increasing intraocular pressure steps was visualized by spectral domain optical coherence tomography (SD-OCT) imaging. Representative images at three pressure steps (10, 15 and 20 mmHg) in three different age groups are shown. In each panel, the upper row shows the OCT image, and the lower row shows the same image in which SC lumen has been identified (blue) by our segmentation software. **A**: young, **B**: middle-aged and **C**: elderly.

The relative change in SC luminal cross-sectional area in response to IOP elevation was calculated for each animal in each of the three age groups (Figure 4A-C). For each eye the slope of the relationship between relative SC luminal area and IOP was then calculated, with larger negative values representing greater narrowing of the SC lumen due to increased IOP. After compiling the slopes for all eyes, we observed an age-dependent increase in the resistance to narrowing of SC as IOP increased (*R*^2^ = 0.55, *p* < 10^−3^, Figure 4D). This observation suggests that TM tissues become structurally stiffer with age.

**Figure 4.**
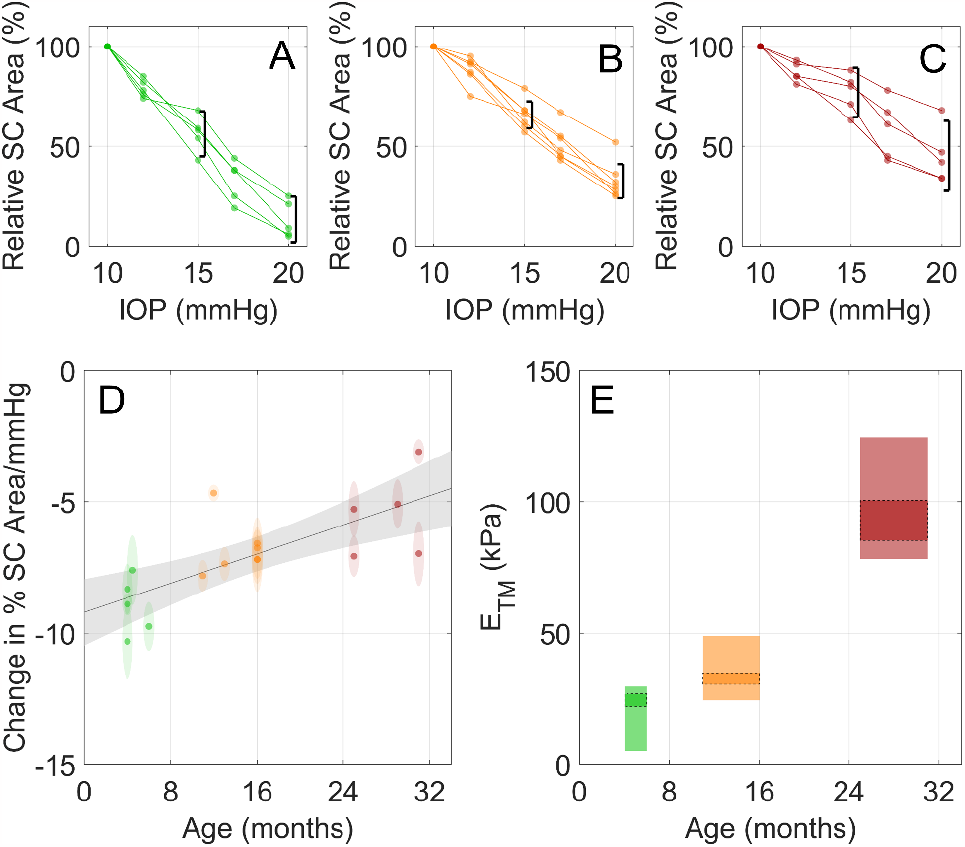
Quantification of reduced SC collapse and trabecular meshwork stiffening with age in mice. SC luminal cross-sectional area was evaluated as a percentage of the baseline (10 mmHg) SC luminal cross-sectional area for young (**A**), middle-aged (**B**) and elderly (**C**) mice, using images such as those shown in Figure 3. Lines connect data points for a given eye. Black brackets indicate 95% confidence intervals on Relative SC Area at 15 and 20 mmHg for all mice in the given age group, which were used as inputs for finite element modelling to determine effective TM stiffness. **D**) The slopes of the lines in panels A, B and C, computed using linear regression for each eye, are seen to be correlated with age (R^2^ = 0.55, p < 0.001). Ellipse heights indicate 95% confidence intervals on the slopes and widths indicate age resolution of 1 month. Line shows the best linear fit with grey shaded region indicating the 95% confidence band. **E**) Trabecular meshwork stiffness was estimated by forward finite element modelling, as described in the text. The vertical limits of the rectangles indicate the range of TM stiffness (Young’s modulus, *E*_*TM*_ ) values that predict SC areas within the 95% confidence of the mean of the experimentally observed SC narrowing, while widths indicate the age range of the experimental data. Rectangles with dashed outlines indicate Young’s modulus values calculated when considering SC collapse as IOP was increased from 10 to 15 mmHg, while those without borders represent values calculated when considering SC collapse as IOP was increased from 10 to 20 mmHg. See Table S1 for plotted values. Colors: green – young, orange – middle-aged, red - elderly.

We next used forward finite element modeling to estimate TM stiffness based on fitting the computed collapse of SC lumen to observed values of relative SC collapse at pressures of 15 and 20 mmHg for each age group (Supplementary Figure 2). We computed a significant increase in the Young’s Modulus of the TM with age; for example, over the IOP range from 10-15 mmHg, we estimated an age-associated increase of Young’s modulus of 28-37% in middle-aged vs. young mice, and of 273-285% in elderly vs. young mice). Interestingly, this stiffening behavior appeared nonlinear, i.e. there was accelerated stiffening with age (Supplemental Table 1, Supplemental Figure 2, Figure 4E).

### Outflow facility and IOP are stable with age

Despite the loss of cells, increased deposition of pigment and increased TM tissue stiffness, IOP was remarkably insensitive to age in mice (*R*^2^ = 0.00, *p* > 0.05, Figure 5A). To evaluate how physical changes in the TM with age affected its function, we used a specially designed system (iPerfusion) to measure the hydraulic conductance of the conventional outflow tissues (outflow facility) in enucleated eyes from mice of different ages [48]. Although we observed a gradual increase in outflow facility with age, the correlation was weak and when corrected for multiple comparisons the slope was statistically different from zero, albeit this was borderline (*R*^2^ = 0.26, *p*_*B*_ < 0.05, Figure 6B). When adjusting for the increase in the filtering area of the outflow pathway with age (due to increased globe size), the correlation between age and adjusted outflow facility was marginally weaker and not statistically different from zero (*R*^2^ = 0.20, *p*_*B*_ > 0.05, Supplemental Figure 3A).

### Age alters the response of anterior chamber structures to IOP

The mechanical responses of anterior segment tissues in *ex vivo* eyes to increasing IOP was also analyzed by calculating the non-linearity in the pressure-flow curves obtained during perfusions of enucleated eyes, characterized by the parameter β [48]. Specifically, larger β values correspond to an increase in outflow facility with increasing pressure and are thought to be associated with more posterior iris/lens movement during cannulation of the anterior chamber (i.e., anterior chamber deepening) [49]. Interestingly, we observed that β decreased in an age-dependent manner, with an average reduction of -0.027 ± [-0.039, -0.015] per month (*R*^2^ = 0.51, *p*_*B*_ < 0.001, Figure 5C). These data indicate that β is effectively zero in elderly eyes. Similarly, a strong inverse correlation was observed between adjusted ocular compliance and age (*R*^2^ = 0.82, *p*_*B*_ < 0.001, Figure 5D), a measure of ocular coat distensibility, indicating that the corneoscleral shell stiffens with age. Even without adjustment for changes in ocular diameter with age, the correlation between ocular compliance and age was still observed (*R*^2^ = 0.56, *p*_*B*_ < 0.001, Supplemental Figure 3B).

**Figure 5.**
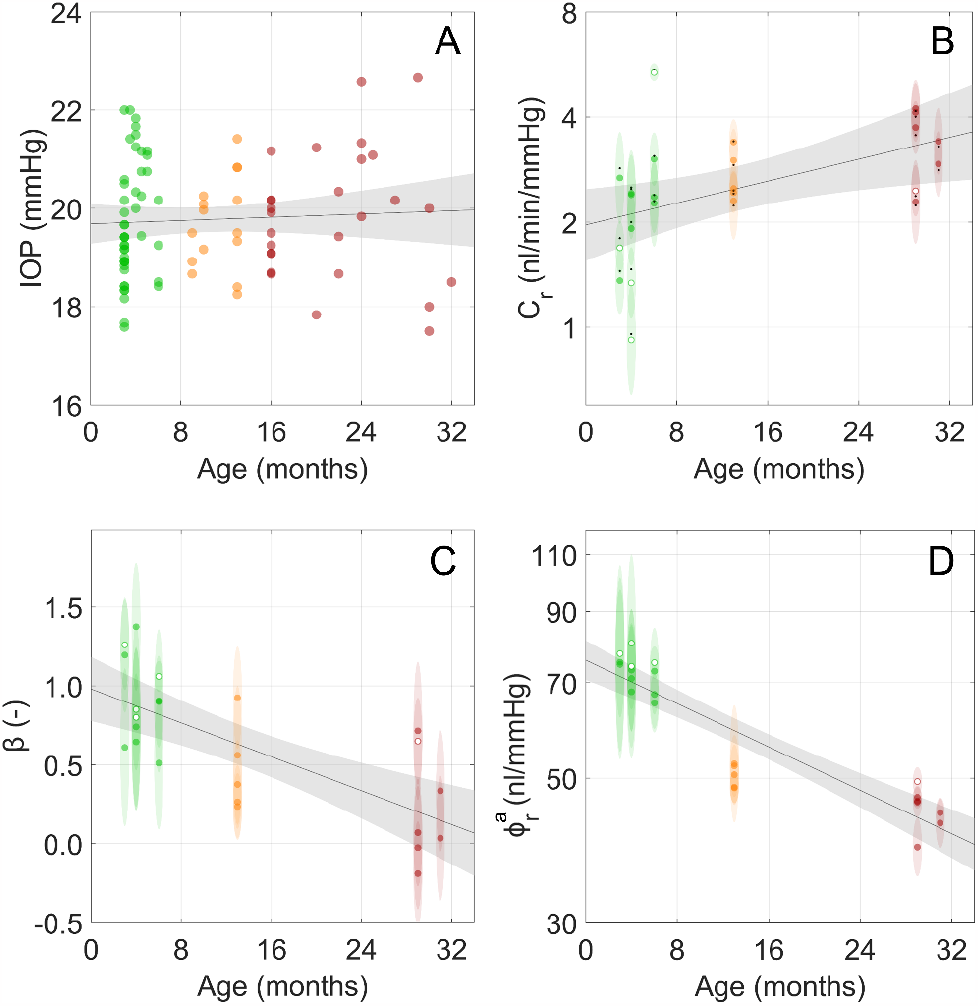
While IOP and outflow facility were relatively insensitive to age, the pressure-dependence of outflow facility (β) and ocular compliance both decreased with age. **A)** IOP was measured by tonometry. No correlation between age and IOP was observed (*R*^2^ = 0.00, *p* > 0.05). **B)** Measured outflow facility showed a weak correlation with age (*R*^2^ = 0.26, *p*_*B*_ < 0.05). Ellipse heights indicate 95% confidence intervals on facility and widths indicate the age resolution of 1 month. Hollow circles come from individual eyes while filled circles are averaged between both eyes from an animal. Black dots in panel B indicate adjusted facility values shown in Supplemental Figure 3A. **C)** A reduction in β, which represents the relative increase in outflow facility as pressure increases, was observed with increasing age (*R*^2^ = 0.51, *p*_*B* < 0.001). Ellipse heights indicate 95% confidence intervals on β and widths indicate age resolution of 1 month. **D)** A strong negative correlation was observed between adjusted ocular compliance, 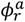 (see text), measured via perfusion and adjusted for eye size to the expected compliance at 3.4 mm diameter, and age (*R*^2^ = 0.82, *p*_*B*_ < 0.001). Lines show best linear fits with grey shaded regions indicating the 95% confidence bands. Colors: green – young, orange – middleaged, red - elderly.

Using OCT imaging, we also monitored iris deformations in response to stepwise increases of IOP (Figure 6). We observed that although the irides of older eyes were more posteriorly located at a low IOP of 10 mmHg vs. in young and middle-aged eyes (Figure 6A), the irides in young eyes were more susceptible to posterior deformation and bowing as IOP increased vs. in older eyes. To quantify this effect, we measured the angle between the iris and cornea for each animal from OCT scans, and for all cases observed an increase in this subtended angle with increasing IOP that appeared reversible when returning to a 10 mmHg baseline IOP (Figure 6B-D). From these observations, we further quantified how the baseline angle (at 10 mmHg) varied with age, and found an average increase of 1.2 [0.7,1.8] °/month (*R*^2^ = 0.64, *p* < 10^−3^, Figure 6E). Further, we observed that the sensitivity to changes in IOP decreased with age and quantified this effect through a constant of proportionality, *ζ*, between the change in angle and the change in pressure from baseline. We observed a weak correlation between *ζ* and age (*R*^2^ = 0.38, *p* < 0.05, Figure 6F), which is consistent with structural stiffening of the iris.

**Figure 6.**
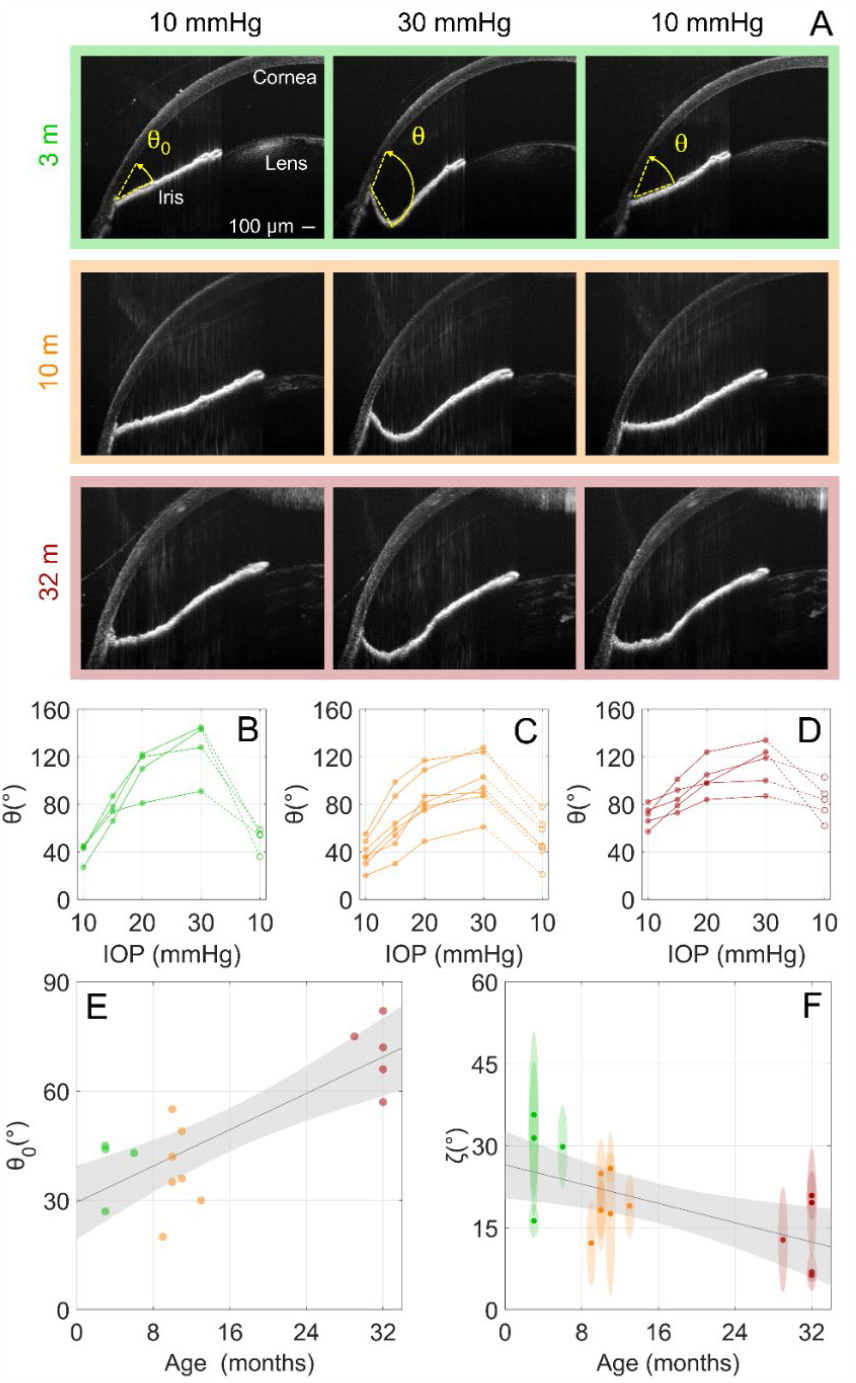
Iris deformations in response to increased IOP decreased with age in mice. **A)** Representative cross-sectional OCT images of mouse anterior segments in young, middle-aged and elderly mice. Images are shown (left to right) at the baseline IOP of 10 mmHg, at the peak IOP of 30 mmHg, and after returning to 10 mmHg. **B-D)** Quantitation of the irido-corneo angles at different IOPs from three age groups of mice (B: young, C: middle-aged and D: elderly). Connecting lines indicate individual eyes, and the dashed lines indicate the reduction in IOP back to the baseline of 10 mmHg. **E)** The irideo-corneal angle at the baseline IOP (10 mmHg, θ_0_) was correlated with age (R^2^ = 0.64, p < 0.001). **F)** The increase in irideo-corneal angle from baseline (θ-θ_0_) was evaluated relative to the logarithm of the increase in IOP from baseline (P-P_0_) with constant of proportionality ζ, which was calculated using regression. ζ represents the sensitivity of the irido-corneo angle to increasing IOP and was found to be negatively correlated with age (R^2^ = 0.38, p < 0.05). Ellipse heights indicate 95% confidence intervals on ζ and widths indicate age resolution of 1 month. Lines show best linear fits with grey shaded regions indicating the 95% confidence bands. Colors: green – young, orange – middle-aged, and red – elderly.

## 4. Discussion

In the present study, we examined a number of age-related changes in the conventional outflow tissues of mice, responsible for regulating IOP. With age, we observed an increase in pigment and pigment-laden cells in the conventional outflow pathway, but a decrease in the total number of cells and increased levels of the senescence marker CDKN1A (p21). These morphological changes were accompanied by alterations in the biomechanical properties of associated ocular tissues, which included stiffening of the TM, decreased pressure-induced deepening of the anterior chamber and decreased ocular compliance. Despite this myriad of age-related changes in the eye, especially in the outflow tissues, outflow facility remained stable or even increased slightly with age to maintain a constant IOP over the life of the mice. Thus, much like humans, the healthy mouse eye appears to adapt to agerelated changes to maintain IOP within a narrow range over a lifetime [50, 51], reinforcing the idea that homeostatic feedback loop(s) regulate IOP [52, 53].

We observed more pigment and fewer cells in TM tissues, especially in the JCT area, with increasing age. These results are strikingly similar to age-related changes in the human TM, which also shows more pigment and fewer cells with age [32, 33, 54, 55]. Significantly, there are even fewer TM cells and more pigment in glaucomatous eyes compared to age-matched controls [32, 54]. These results suggest a reduced capacity of TM cells, and possibly resident macrophages, to process pigment that is released by the iris and other pigmented ocular epithelia. The TM functions as a biological filter, and thus a primary role for the TM cells is to neutralize oxidants and phagocytose cellular debris carried by the flow of aqueous humor into the TM [56-58]. TM cells are assisted in this function by resident macrophages. Consistent with this understanding, we observed more pigment-laden cells with increasing age. The remaining TM cells in older eyes showed signs of senescence (i.e., increased CDKN1A expression), possibly due to increased phagocytic and oxidative loads [59]. CDKN1A is a cyclin-dependent kinase inhibitor that is a major target of p53 activity and thus is associated with linking DNA damage to cell cycle arrest [60].

Accompanying these morphological alterations in the TM with age, the biomechanical properties of ocular tissues, including the TM, also changed. Using our novel combination of OCT imaging of outflow tissues subjected to varying IOP and finite element modeling, we observed significant stiffening of the TM with age. Values for Young’s modulus obtained for the TM were consistent with our previous reports [38, 61]. For example, over the physiological IOP range 10-15 mmHg, we estimated TM stiffnesses of 22-27 kPa in young mice in the present study, which was similar to the 25-29 kPa estimated previously in a young cohort. While age-induced stiffening of the TM has not been examined previously in human eyes, the TM has been shown to be stiffer in glaucomatous eyes compared to age-matched controls [44, 45, 62, 63]. Such changes in biomechanical properties can be induced in non-glaucomatous human TM cells or in the TM of young mice by prolonged glucocorticoid treatment [39]. Using our new finite element model that accounts for physical changes in the eye with age, we estimated a TM stiffness of 85-101 kPa over the physiological IOP range 10-15 mmHg for our elderly cohort in the present study, which is greater than the estimated stiffness of 69 kPa for a younger cohort of mice with steroid-induced ocular hypertension [38].

Taken together, these data suggest important differences between normal age-associated changes and pathological changes to the TM. More specifically, increased TM tissue stiffness positively correlates with a *decrease* in outflow facility in young mice under pathological conditions [45], while in the present study, we saw that both TM stiffness and outflow facility remained stable with age. This apparent paradox may be due to differences in disease versus normal aging, both of which result in TM stiffening but by different biological mechanisms. For example, there was an increase in basal laminar deposits below the inner wall of SC in young mice experiencing glucocorticoid-induced ocular hypertension and TM stiffening [39], whereas in the present study we observed the normal “patchy” basal laminar materials below the inner wall of SC in elderly mice with a stiffer TM.

Part of the observed minor increase in facility (Figure 5B and Supplementary Figures 1C and 3A) can be attributed to age-related morphological changes such as increased globe size, and thus a corresponding increase in TM cross-sectional area with age [64]. After adjusting for TM area, the correlation coefficient became statistically indifferent to zero, implying no change in conductivity of the TM/SC. The small increase in un-adjusted facility may be a compensatory effect associated with decreased unconventional outflow with age, as seen in non-human primates. This would result in a shunting of flow from the unconventional to the conventional outflow tract [65], which in turn would require an increase in facility to maintain a desired IOP. Such an increase is consistent with two previous studies that examined outflow facility in aging mice; each reported an increase, or a trend towards an increase, in outflow facility in 3 of 4 strains studied, including C57 mice [66, 67].

In addition to increased TM tissue stiffness, we observed other age-related changes in ocular tissue biomechanical properties. For example, we monitored posterior iridial bowing due to increasing IOP. Like TM tissues, the pressure-induced posterior displacement of the iris decreased with age, indicating increased iridial structural stiffness. These results are consistent with a decreased β that was observed in older eyes during ocular perfusions in the present study. β is the constant of proportionality between relative changes in facility and relative changes in pressure [49], and a smaller β is consistent with less posterior displacement of the iris and lens as IOP increases, likely due to increased ocular tissue stiffness. Lastly, we observed decreased ocular compliance with age. Even when adjusted for increased globe size with age, compliance was still significantly lower in older eyes. These three indicators of changes in ocular tissue properties with age are consistent with an abundant literature demonstrating that age decreases tissue elasticity and increases tissue stiffness in many other tissues such as skin, lungs, and blood vessels [37].

In summary, we provide the first comprehensive study reporting the effects of age on the properties and function of conventional outflow tissues. As the primary determinant of intraocular pressure (and thus playing a critical role in ocular hypertension and glaucoma), it is important to understand how the function and morphology of outflow tissues change with age. The changes we describe here provide a physiological backdrop for susceptibility to genetic or environmental insults that result in outflow dysfunction/loss of IOP homeostasis, elevated IOP and glaucoma.

## Author Contributions

GL, JvBS, BNS, NSFG, MRBF, CBR, AJF, SF, CRE and WDS contributed to the study’s conception and design. GL, JcVS, KC, MLD, AJF, SF, CRE and WDS conceived and designed the experiments; FL, VNS, AW, KC, MLD, SC, TW, MK, performed the experiments; GL, JvBS, BNS, NSFG, MRBF, KC, MLD, SC, AJF, MK, SF, CRE and WDS analyzed the data; GL, JvBS, CRE and WDS wrote, reviewed and edited the manuscript. All authors read and approved the final manuscript.

## Acknowledgements

We gratefully acknowledge support from The BrightFocus Foundation (postdoctoral fellowship G2021005F, BNS and Special Opportunity Award, WDS, CRE and JvBS), NIH (K99EY035360 (BNS), T32EY007092 (NSFG), R01EY030871 (AJF), P30EY006360, P30EY005722, R01EY031748 (CBR), R01EY031710 (CRE and WDS), R01EY030124 (WDS) the Alfred P. Sloan Foundation G-2019-11435 (NSFG), the Georgia Research Alliance (CRE), and Department of Veterans Affairs Rehab R&D Service Career Development Awards (AJF; CDA-2; RX002342).

## Data Availability Statement

Data available on request from the authors

