## Supplementary files for "Aging and intraocular pressure homeostasis in mice"

### Supplementary Figures and Table

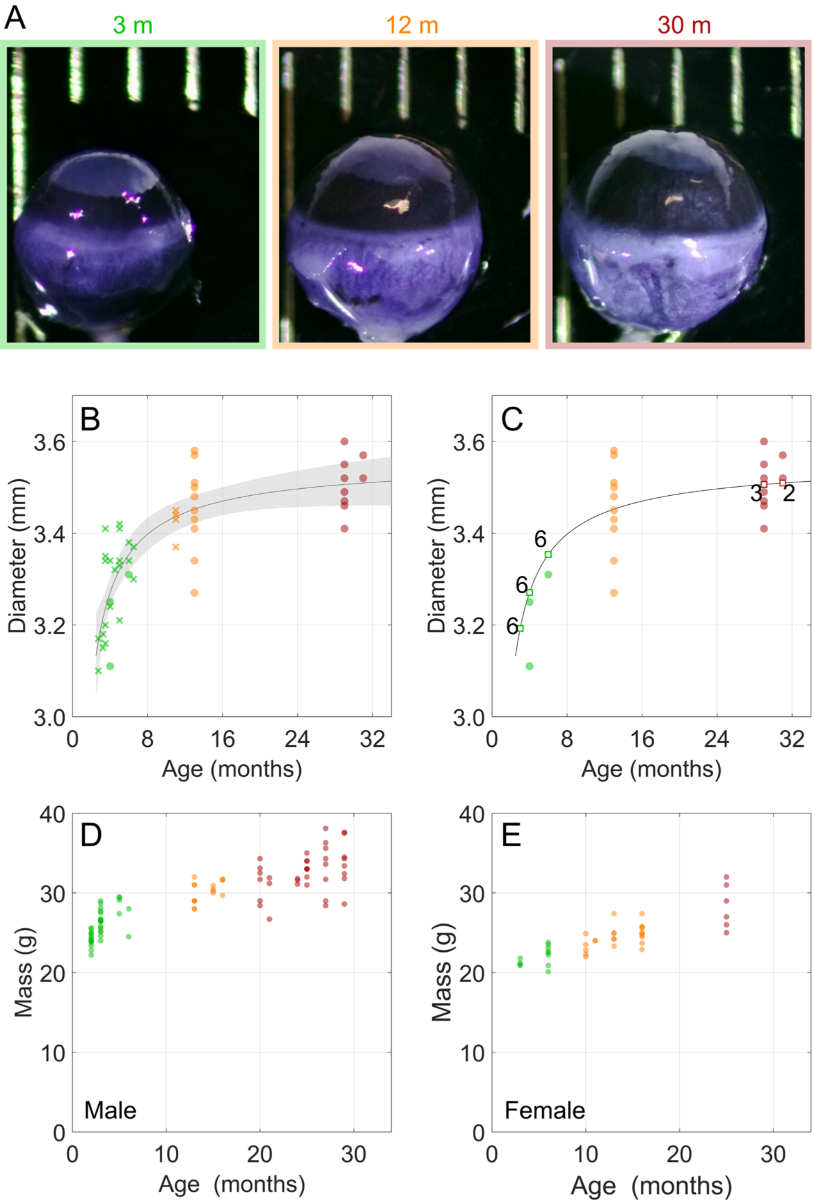


**Supplemental Figure 1. Age-related changes in eye size and mass of mice. A)** Representative images of eyes from young, middle-aged and elderly groups. **B)** Ocular size (diameter) increased with age. Circles indicate data for which both aqueous humor dynamics (AHD) parameters and ocular diameter were available, while crosses indicate data points for which only diameter data was available. The curve shows best fit to the empirical equation $d = a_1 \, Age^{\, a_2} + a_3$, where $d$ is ocular diameter, the $a_i$ are fitting coefficients, and the grey shaded area indicates 95% confidence bands. **C)** Diameter values used for adjustment of facility and ocular compliance. The plotted curve is identical to that shown in panel A. Circles indicate data for which both AHD parameters and diameter were available, and squares indicate data points for which diameter was interpolated based on the fit shown in panel A. Numbers indicate how many eyes were interpolated to that diameter. Body mass for male (**D**) and female (**E**) mice increased with age. Colors: green – young, orange – middle-aged, and red - elderly.

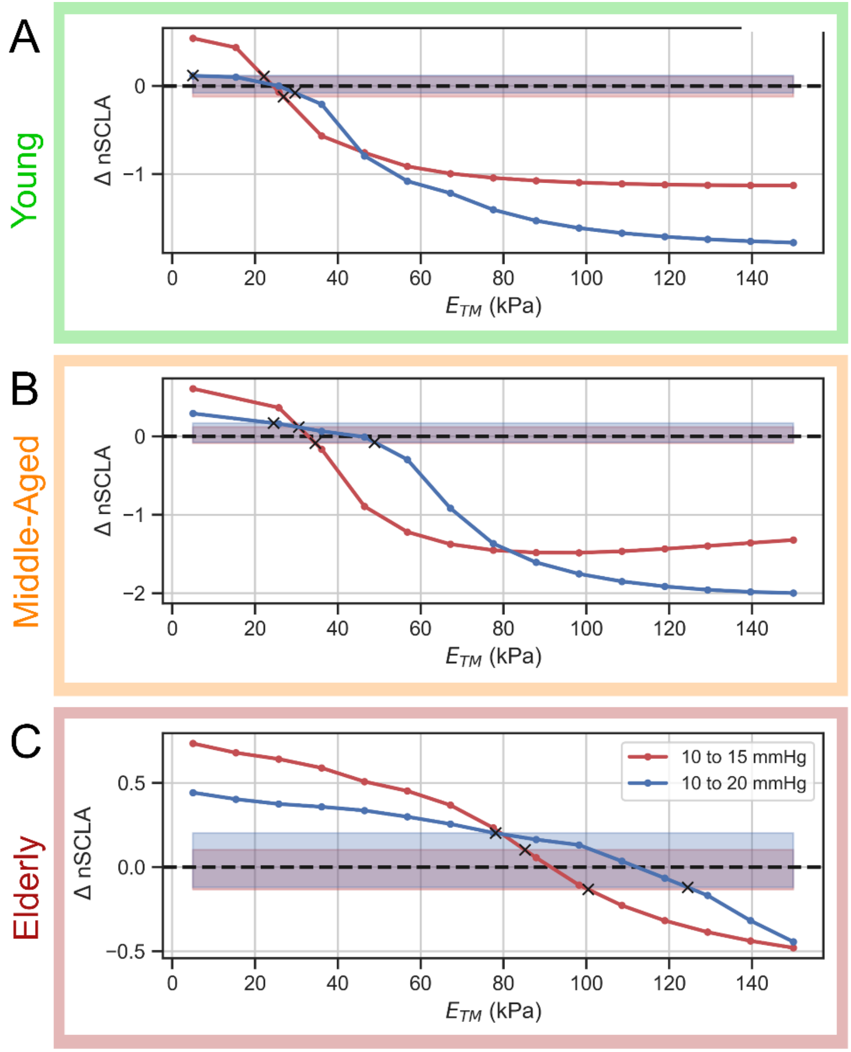


**Supplemental Figure 2. Forward finite element simulations were used to estimate Young's modulus of the TM, $\boldsymbol{E_{TM}}$, for young (A), middle-aged (B) and elderly (C) eyes.** The plotted quantity, $\boldsymbol{\Delta nSCLA}$, is defined in the text and is a measure of how closely the computed collapse of SC lumen agrees with the experimentally observed collapse as IOP is increased from 10 to 15 mmHg (red), or from 10 to 20 mmHg (blue). The blue and red shaded horizontal bands, which overlap to some extent, are a measure of the 95% confidence intervals on the experimental data (see black brackets in Figure 4A-C). The intersections of the $\boldsymbol{\Delta nSCLA}$ curves with these confidence bands are denoted by "x", which correspond to the upper and lower limits of $\boldsymbol{E_{TM}}$ values for which the computed and experimental measures of SC lumen collapse are consistent; it is these limits that are plotted in Figure 5E.

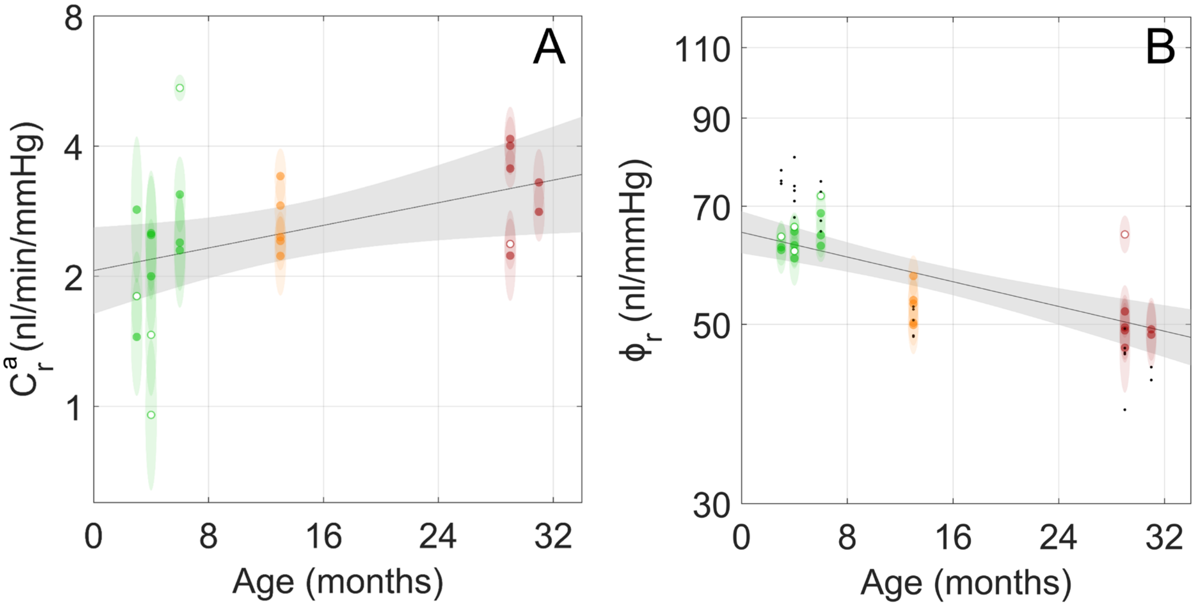


**Supplemental Figure 3. Adjustments to facility and ocular compliance to account for eye size did not affect overall conclusions. A)** Outflow facility measured via iPerfusion and adjusted for eye size to the expected facility at 3.4 mm, $C_{r,i}^{a}$ (see text for details). The weak correlation between the logarithm of adjusted facility and age was not statistically different from zero ($R^2$ = 0.20, $p_B > 0.05$). **B)** Ocular compliance prior to adjustment. A moderate dependence on age was observed ($R^2$ = 0.56, $p_B < 0.001$). Dots show adjusted values (Figure 8B). Ellipse heights indicate 95% confidence intervals on facility and ocular compliance, while widths indicate age resolution of 1 month. Hollow circles come from individual eyes while filled circles are values averaged between both eyes of an animal. Colors: green – young, orange – middle-aged, and red – old.

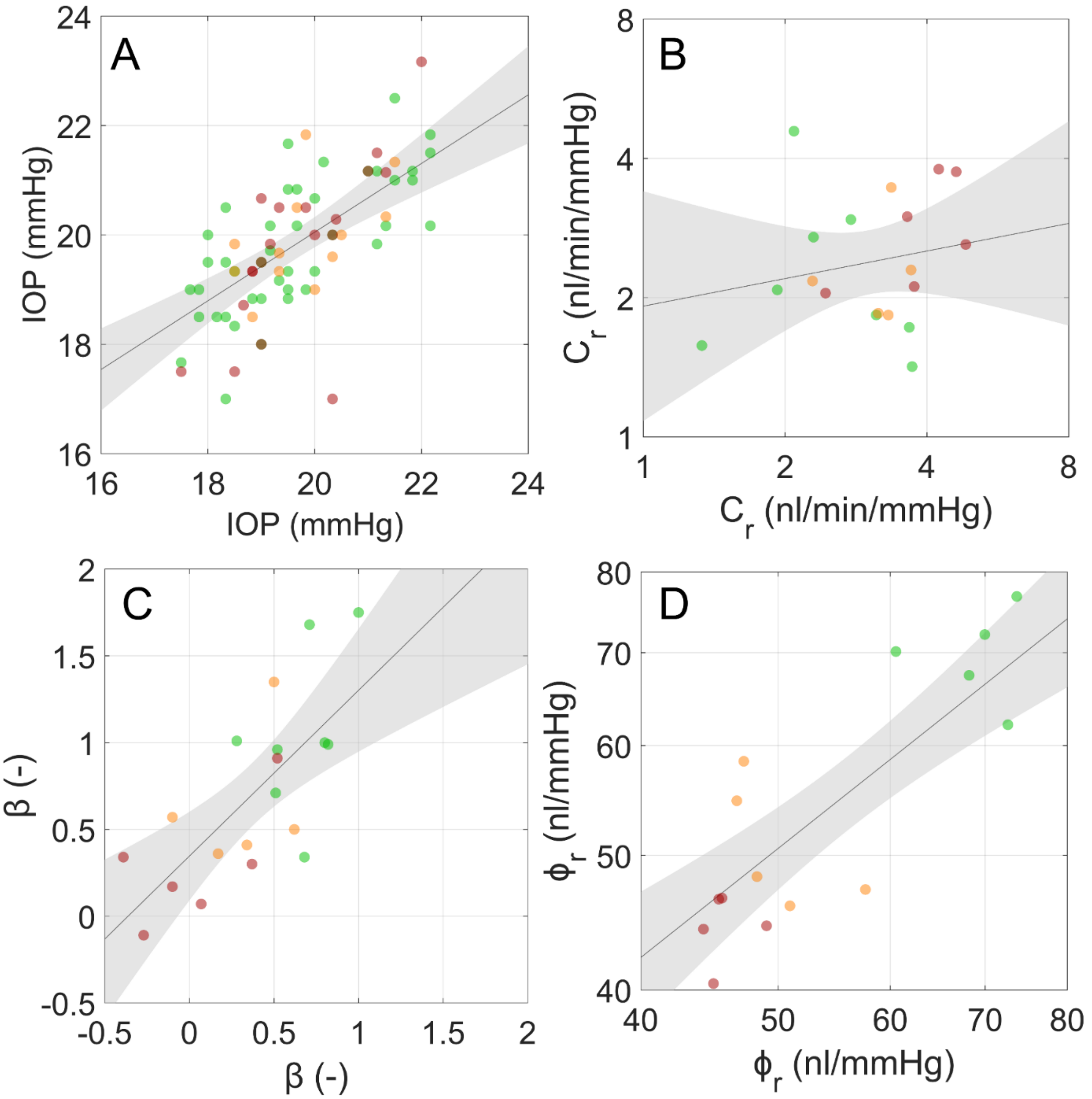


**Supplemental Figure 4. Analysis of correlation between paired eyes of mice for aqueous humor dynamics (AHD) parameters.** All AHD parameters except facility were correlated between two eyes of a given animal ($R^2 > 0.5$, $p_B < 0.01$); hence, all paired data were averaged and treated on a per animal basis in statistical analysis. Each data point on each graph arises from paired eye measurements on a single mouse, showing: **A)** IOP, **B)** the logarithm of the eye size-adjusted reference outflow facility, **C)** beta, and **D)** the logarithm of the eye size-adjusted reference ocular compliance. Colors: green – young, orange – middle-aged, and red – elderly.

**Supplementary Table 1:** Computed ranges for Young's modulus of the trabecular meshwork, $\boldsymbol{E_{TM}}$, as plotted in Figure 4E.

| **Age** | **IOP range (mmHg)** | $\boldsymbol{E_{TM}}$ **(kPa)** |
|---|---|---|
| Young | 10-15 | 22.1-26.9 |
| | 10-20 | 5.0-29.6 |
| Middle-aged | 10-15 | 30.5-34.6 |
| | 10-20 | 24.5-48.8 |
| Elderly | 10-15 | 85.2-100.5 |
| | 10-20 | 78.1-125.0 |
